# ECCsplorer: a pipeline to detect extrachromosomal circular DNA (eccDNA) from next-generation sequencing data

**DOI:** 10.1101/2021.06.08.447410

**Authors:** Ludwig Mann, Kathrin M. Seibt, Beatrice Weber, Tony Heitkam

## Abstract

**Motivation:** Extrachromosomal circular DNAs (eccDNAs) are ring-like DNA structures physically separated from the chromosomes with 100 bp to several megabasepairs in size. Apart from carrying tandemly repeated DNA, eccDNAs may also harbor extra copies of genes or recently activated transposable elements. As eccDNAs occur in all eukaryotes investigated so far and likely play roles in stress, cancer, and aging, they have been prime targets in recent research – with their investigation limited by the scarcity of computational tools.

**Results:** Here, we present the ECCsplorer, a bioinformatics pipeline to detect eccDNAs in any kind of organism or tissue using next-generation sequencing techniques. Following Illumina-sequencing of amplified circular DNA (circSeq), the ECCsplorer enables an easy and automated discovery of eccDNA candidates. The data analysis encompasses two major procedures: First, read mapping to the reference genome allows the detection of informative read distributions including high coverage, discordant mapping, and split reads. Second, reference-free comparison of read clusters from amplified eccDNA against control sample data reveals specifically enriched DNA circles. Both software parts can be run separately or jointly, depending on the individual aim or data availability. To illustrate the wide applicability of our approach, we analyzed semi-artificial and published circSeq data from the model organisms *H. sapiens* and *A. thaliana,* and generated circSeq reads from the non-model crop *B. vulgaris.* We clearly identified eccDNA candidates from all datasets, with and without reference genomes. The ECCsplorer pipeline specifically detected mitochondrial mini-circles and retrotransposon activation, showcasing the ECCsplorer’s sensitivity and specificity. The derived eccDNA targets are valuable for a wide range of downstream investigations – from analysis of cancer-related eccDNAs over organelle genomics to identification of active transposable elements.

**Availability and implementation:** The ECCsplorer pipeline is available on GitHub at https://github.com/crimBubble/ECCsplorer under the GNU license.

**Contact:** Tony Heitkam

**Supplementary information:** Supplementary data are available online.

## 1 Introduction

Although first described over 50 years ago, the physiological role of most extrachromosomal circular DNAs (eccDNAs) remains debated (Hotta and Bassel, 1965; Liao *et al*., 2020; Møller *et al*., 2019; Molin *et al*., 2020). Nevertheless, eccDNAs are often associated with many different biological roles and processes. In animals and human they have been reported as drivers for aging (Gaubatz and Flores, 1990; Sinclair and Guarente, 1997) and cancer(Turner *et al*., 2017; Kumar *et al*., 2017; Paulsen *et al*., 2018), whereas in plants they also play a role in gradual development of glyphosate resistance(Benoit, 2020). Many of these physiological consequences can be traced to extra gene copies residing extrachromosomally within the DNA circles. Beside including genes, eccDNAs often consist of repetitive DNAs such as tandem repeats (satellite DNAs) or (active) transposable elements (Flavell and Ish-Horowicz, 1981; Lanciano *et al*., 2017; Møller *et al*., 2019; Diaz-Lara *et al*., 2016). Thus, due to their relationship with transposable elements and genes, eccDNAs may also be involved in transcriptional regulation, e.g. mediated by small RNAs (Paulsen *et al.*, 2019).

Although few functions of eccDNAs are known so far, there is a growing interest from medicine (e.g. early disease detection or therapy) and economy (e.g. new traits for crop plants through transposon activity or gene accumulation). Therefore, many emerging studies focus on eccDNAs in model and non-model organisms. This increasing interest comes along with new challenges for eccDNA detection, both experimentally and bioinformatically. Most recent approaches rely on next-generation sequencing after experimental enrichment of eccDNA (Lanciano *et al*., 2017; Møller *et al*., 2019; Diaz-Lara *et al*., 2016; Shibata *et al*., 2012). To enrich double-stranded circular DNA, phi29 polymerases are usually used for a random rolling circle amplification (rRCA) after linear was removed by an exonuclease. Although single long-read approaches are described (Mehta *et al*., 2020), short-read techniques are widely common to generate so called circSeq data (in special cases also referred to as mobilome-Seq data, e.g. Lanciano *et al*., 2021). Computationally, new eccDNA candidates are currently identified based on typical read signatures, commonly detected by the mapping against a reference genome (Lanciano *et al*., 2017; Prada-Luengo *et al*., 2019). These characteristic signatures include split and discordant mapping reads, which result when mapping a circular sequence against a linear reference. Based on analysis of those reads, the most likely eccDNAs are commonly called. Yet, these properties are not unique to eccDNAs – repetitive DNAs, structural variants and assembly errors in the reference yield a similar read spectrum – thus, possibly resulting in many false positives.

To date, only very few software solutions are available to analyze the growing amount of circSeq data, and currently no solution is able to address the aforementioned challenges in a single approach. In addition, there is no software solution, yet, to analyze eccDNAs from short reads if a reference genome sequence is lacking.

Here, we present the ECCsplorer pipeline – a bioinformatics approach to specifically detect candidates with high confidence from eccDNA-enriched sequence datasets. Our pipeline is modular to guarantee maximal flexibility and reliability and represents a reproducible way for the detection of eccDNAs in both model and non-model organisms. It supports standard input formats (un-/compressed FASTQ and FASTA) as provided by most sequencing services and is available at https://github.com/crimBubble/ECCsplorer. We demonstrate its wide applicability using a combination of real and simulated sequence reads from three organisms, including model- and non-model organisms from two kingdoms. This allows us to showcase the ECCsplorer pipeline’s specificity and sensitivity by targeting a broad range of eccDNA candidates: partial gene copies *(Homo sapiens)* active transposable elements *(Arabidopsis thaliana*), and mitochondrial minicircles (*Beta vulgaris*).

## 2 Material and methods

### 2.1 Overview

The ECCsplorer pipeline (Fig. 1a) is modular and implemented in Python 3 and partly in R (R Core Team, 2013). The integration of the statistical programming language R provides the capability of performance-enhancing as well as a user-friendly output for publication-ready figures. The ECCsplorer pipeline is allowing three input configurations (`running modes: all/map/clu`) to answer a variety of scientific questions (Fig. 1a). First, circSeq data from rRCA enriched DNA is mandatory to start the pipeline. Second, either a control dataset (`mode: clu`) or a reference genome sequence (`mode: map`) is necessary. As control data a variety of sequencing data can be provided for use (e.g. rRCA enriched data from a different ecotype or tissue, non-enriched data, or WGS data). Using both control data and a reference genome sequence, the ECCplorer yields the most reliable results and represents the third input configuration (`mode: all`). Additionally, a user-defined file of reference sequences (containing e.g. known repeats, genes, etc.) might be provided for candidate annotation.

**Figure 1:**
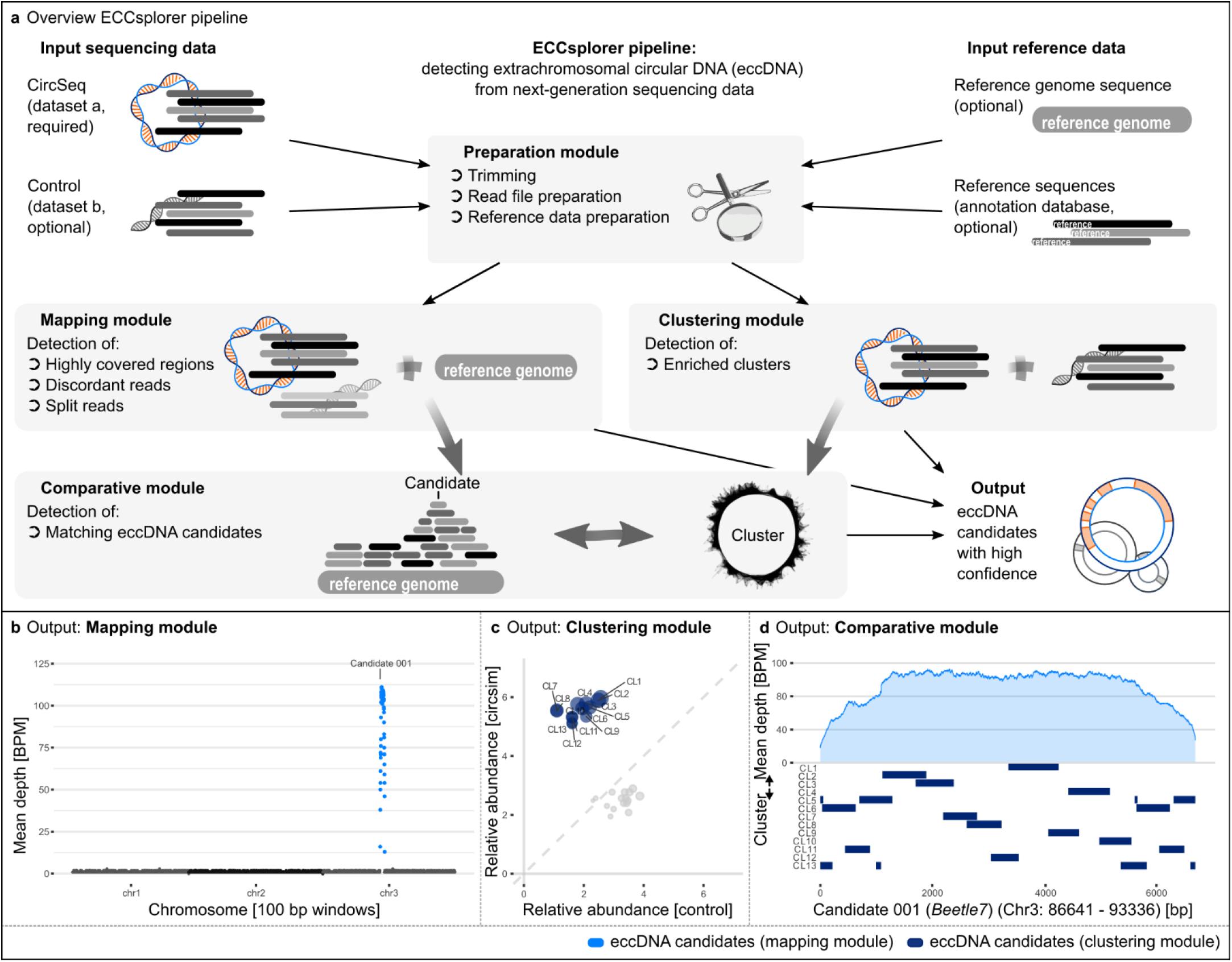
Overview of the ECCsplorer pipeline (**a**) and it’s output (**b-c**). (**a**) Our pipeline consists of four main modules: one for the data preparation module and three for the analysis modules. The execution of the different modules depends on the given input data. Thick grey arrows represent multi-step bioinformatics computation processes, thin black arrows show data transfer. Semi-artificial circSeq and control data were simulated from a circularized retrotransposon and a random reference sequence to illustrate a typical pipeline output. (**b**) Mapping against the reference sequence revealed an eccDNA candidate region. (**c**) The comparative read clustering identified 13 clusters containing potential eccDNA candidates. (**d**) The comparative module links the retrotransposon candidate region from the mapping module to all 13 clusters from the clustering module.

The ECCsplorer workflow can directly use raw sequence data as input and covers the steps for read file and reference preparation. The so-called, preparation module includes read trimming (using Trimmomatic; Bolger *et al*., 2014) and sampling as well as file conversion and indexing of the reference sequence.

The main part of the pipeline controls the analysis and consists of three additional modules: the mapping module, the clustering module and the comparative module. Depending on the given input data either the mapping module, the clustering module, or both are started (Fig. 1a):

- The mapping module maps the circSeq data to a reference sequence (e.g. genome assembly) with the tool segemehl (Hoffmann *et al*., 2014). Use of an additional control dataset will improve the module’s results, but is not obligatory for the mapping itself.
- The clustering module is only executed, if control data are provided in addition to the circSeq dataset. This module conducts a comparative read clustering analysis with the RepeatExplorer2 pipeline (Novák *et al*., 2010, 2020).
- If all three input datasets are available, the results of both modules will be combined by the comparative module using a similarity search with the BLAST+ package (Camacho *et al.*, 2009).

The ECCsplorer pipeline will output the resulting eccDNA candidates for each analysis module summarized in up to three HTML files including publication-ready figures. In addition, the R script underlying all visualizations is available to the user for customization.

### 2.2 The preparation module

The preparation module processes all input data and checks for the required software packages, libraries and third-party tools. If the recommended trimming option is enabled, Trimmomatic (Bolger *et al*., 2014) is used to ensure high quality input for the ECCsplorer pipeline. Then, the read files are converted from FASTQ to FASTA if necessary using either seqtk (Li *et al*., 2013) (if pre-installed) or the converter from the SeqIO (Cock *et al*., 2009) python package. In the next steps the read files are prepared for the read clustering with RepeatExplorer2 (Novák *et al*., 2010, 2020). In short, read files are sub-sampled, interlaced, concatenated, read names are modified with pre- and suffix, and the read lengths are equalized over all reads and files. For the mapping, the reference genome file is indexed. To enable the annotation of detected eccDNA candidates, a BLAST+ database is created, later referred to as annotation database. By default, this annotation database corresponds to the RepeatExplorer2’s internal database and can be extended by a user-defined FASTA file of reference sequences (containing e.g. known repeats, genes, etc.). The user-extended annotation database will be available to the mapping/comparative module. After the data preparation, the analysis modules are started.

### 2.3 The mapping module

The mapping module maps circSeq and control reads (if provided) against a reference sequence and identifies eccDNA candidate regions. For mapping, we use the segemehl (Hoffmann *et al*., 2014) algorithm that was specifically developed for split and circular mappings and is consequently well-suited for detecting eccDNAs. The mapping relies on a reference genome sequence and enables the detection of eccDNA candidate regions. To achieve this, the ECCsplorer relies on the analysis of split reads, discordantly mapping reads, and local high coverages in the following way: Structurally deviating regions flanked by at least five split reads are determined using the haarz (Hoffmann *et al*., 2014) algorithm included within the segemehl tool. Afterwards, regions containing at least one pair of discordantly mapping reads are identified based on the corresponding SAM-flags using SAMtools (Li *et al*., 2009) and BEDtools (Quinlan, 2014). Regions with a significantly high coverage, caused by the experimental enrichment of the eccDNA, are detected using the peak finder algorithm of SciPy (Virtanen *et al*., 2020) with a minimum peak prominence value of one. Finally, all three files with candidate regions are compared for matching or overlapping regions using BEDtools (Quinlan, 2014). The boundaries of the final eccDNA candidate regions are primarily based on the split read regions. Regions meeting all three criteria are considered as candidate regions with high confidence, whereas those complying with any two out of the three criteria are marked as candidate regions with low confidence. Regions with low confidence are only considered in further analysis, if regions with high confidence are absent.

After the final eccDNA candidate regions are determined, a similarity search with BLASTn (Camacho *et al*., 2009) is performed against the annotation database that was previously built. In addition, the candidates’ enrichment scores are calculated using normalized mapping coverages. All coverages are normalized to bases per million bases (BPM), which allow a direct comparison of read inputs of different read lengths. The enrichment score reflects the enrichment by rRCA of a certain eccDNA candidate and is used to rank candidates by their estimated abundance.

### 2.4 The clustering module

For the clustering module, we assume that the presence of eccDNAs leads to the overrepresentation of the encoded sequences. This is additionally enhanced by the experimental amplification. To determine the over-representation of eccDNA sequences, this module performs a comparative read clustering using RepeatExplorer2 (Novák *et al*., 2010, 2020) and requires circSeq and control data as input. With RepeatExplorer2, multiple datasets are comparatively clustered and their repetitive fractions are detected, classified, and quantified. Due to the experimental rRCA, all eccDNAs in the circSeq data exhibit repetitive characteristics, although they might not have originated from true repetitive elements. In contrast to the read mapping the clustering approach is not dependent on a reference genome sequence and less susceptible to background noise. For example, non-circular genomic DNA might still be present in the sample, but in a much lower copy number and will, thus, be filtered by this approach. The key outputs of the RepeatExplorer2 (Novák *et al*., 2010, 2020) utilized within the ECCsplorer pipeline are the cluster-derived contigs and each cluster’s comparative read counts (read count in a cluster per dataset). Clusters with a circSeq read count proportion of more than 80 % are selected for further analysis.

### 2.5 The comparative module

If both the clustering and the mapping modules are executed, the detected eccDNA candidates are further analyzed within the comparative module. For this, contigs of candidate read clusters serve as query for a similarity search with BLASTn (Camacho *et al*., 2009) against the candidate regions detected by the mapping module. Candidates detected by both approaches are considered eccDNA candidates supported with very high confidence and are reported in a conclusive summary.

### 2.6 circSeq data generation and running of the ECCsplorer pipeline

The methods for the generation of circSeq data as well as the explicit commands used in the case studies are available in the supplementary information online (Supp. Info. 1).

### 2.7 Data availability

The semi-artificial test data is available on the ECCsplorers GitHub page (https://github.com/crimBubble/ECCsplorer/testdata). The datasets together with the accession codes are as follows: circSeq (mobilome-Seq) *A. thaliana* (epi12) (Lanciano *et al*., 2017), accession no. ERR1830501; circSeq (mobilome-Seq) *A. thaliana* (WT) (Lanciano *et al*., 2017), accession no. ERR1830499; circSeq *H. sapiens* muscle tissue (Møller *et al*., 2018), accession no. SRR6315430; circSeq *B. vulgaris* KWS2320 (inflorescences), accession no. ERR6004146; WGA *B. vulgaris* KWS2320 (Dohm *et al*., 2014), accession no. SRR869631. Source data for plots shown are included with this paper’s supplementary information (or included in the original articles).

### 2.8 Code availability

All source code for the ECCsplorer pipeline is available at https://github.com/crimBubble/ECCsplorer under the GNU license.

## 3 Results and discussion

### 3.1 Case studies demonstrating the ECCsplorer’s functionality

To evaluate and demonstrate the ECCsplorer’s functionality, we first outlined the results using simulated enrichment of the eccDNA fraction (Fig. 1b-d). Next, we reanalyzed publicly available data of the model organisms *A. thaliana* (Fig. 2a-c) and *H. sapiens* (Fig. 2d), as well as newly generated circSeq data for the non-model crop *B. vulgaris* (Fig. 2e). The three datasets illustrate three different research questions, three different input configurations, and a variety of different eccDNA candidates; thus together fully demonstrate the ECCsplorer pipeline’s wide applicability and functionality.

**Figure 2:**
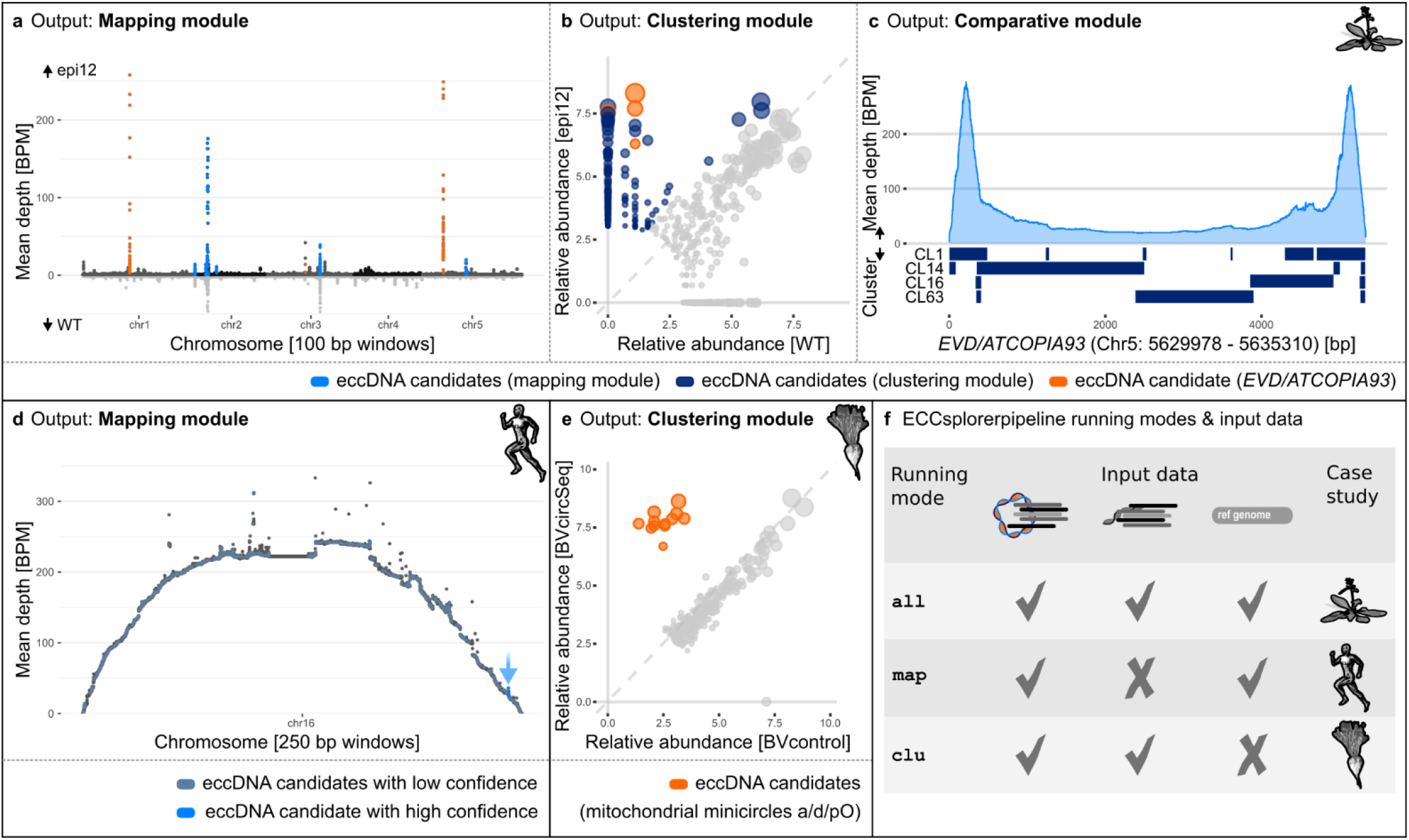
Outputs of the ECCsplorer pipeline for different case studies representing the three running modes and input data configurations. (**a-c**) An ECCsplorer run using *A. thaliana* circSeq (epi12) and control data (WT) (Lanciano *et al*., 2017) detected *EVD/ATCOPIA93* as key eccDNA candidate (orange). (**a**) A mapping against the reference genome sequence (TAIR10) revealed eight eccDNA candidate regions. These were identified by applying three criteria: the region is flanked by split reads (at least five), contains discordant mapping reads (at least one pair), and shows a high coverage compared to its neighboring regions (peak prominence of at least one). (**b**) The comparative read clustering identified 41.04 % of the clustered epi12 reads as potential eccDNA candidates. (**c**) To report eccDNA candidates with high confidence the outputs of the mapping and clustering strategies are compared by the ECCsplorer pipeline. The comparative module links the *EVD/ATCOPIA93* candidate region from the mapping module to four clusters from the clustering module. (**d**) An ECCsplorer run using *H. sapiens* circSeq data (Møller *et al*., 2018) detected one eccDNA candidate region containing genic content (light blue) and many candidate region with low confidence (grey blue). (**e**) ECCsplorer analysis using *B. vulgaris* circSeq data and available sequenceing data (Dohm *et al*., 2014) as control data detected multiple candidate clusters that were in the following annotated as mitchondrial minicircles a/d/pO (orange). (**f**) Overview on the running modes, their required input data and the corresponding case studies.

#### 3.1.1 Case study I: proof of concept using simulated data

To illustrate the ECCsplorer pipeline functionality, we applied it to simulated semi-artificial data (Fig. 1b-d). For this, three regions of the *B. vulgaris* reference genome sequence (Dohm *et al*., 2014) were randomly selected to represent chromosomes with a total length of about 0.6 Mb, including a copy of the well-described long terminal repeat (LTR) retrotransposon *Beetle7* (Weber *et al*., 2013). A potential enrichment of the 6,695 bp long *Beetle7* circular DNA was simulated by introducing a total nine copies of this retrotransposon in tandem arrangement (including solo-LTR and LTR-LTR junctions). To simulate the short-read data, the tool dwgsim (github.com/nh13/DWGSIM) was used. As circSeq data 50,000 paired-end (PE) reads with a length of 200 bp each were simulated using the reference sequence and the tandemly arranged retrotransposon. For the control data the same amount of 200 bp PE reads was generated from the reference only. The reference itself served as reference genome sequence. In addition, the reference sequences of *Beetle1* to *Beetle7* (Weber *et al*., 2013) were used as annotation database.

After trimming, 46,950 PE reads of the circSeq dataset and 46,754 PE reads of the control dataset were piped into the subsequent bioinformatics steps, respectively. Of those reads 12,000 PE reads (6,000 per dataset) were processed further for the clustering analysis, which corresponds to a 4× coverage of the reference. The results from the mapping module as well as those from the clustering module yielded *Beetle7* as the only eccDNA candidate with high confidence. The mapping module reported one candidate region on the artificial reference chromosome 3 with a length of 6,695 bp, an enrichment score of 46.5, and a BLAST+ annotation as *Beetle7,* respectively (Fig. 1b). The clustering module reported 13 candidate clusters (containing 3,475 reads in total, 3,399 reads from the circSeq data and 76 from the control data) adding up to one circular supercluster (Fig. 1c, Supp. Tab. 2). The observed split of the eccDNA candidate into multiple clusters is relatively common after read clustering with RepeatExplorer2. Based on paired end read information, the clusters are commonly linked into superclusters. The comparative module reported that all 13 candidate clusters showed sequence similarities to the eccDNA candidate region (Fig. 1d). Four of those candidate clusters the clusters (namely 5, 6, 11, and 13) cover the circular break-point.

Taken together, the ECCsplorer pipeline successfully retrieved the artificial eccDNA candidates and revealed no other false-positives.

#### 3.1.2 Case study II: identification of active retrotransposons using circSeq data of *A. thaliana*

To demonstrate the ECCsplorer’s functionality to detect the activation of LTR retrotransposons, we re-analyzed public data from *A. thaliana* (Lanciano *et al*., 2017). As a reference genome and both circSeq and control data are available, all ECCsplorer modules were used (`running mode: all`). Lanciano *et. al* amplified eccDNAs of epigenetically impaired *A. thaliana* plants and used the wild-type as control. We expected that our pipeline shows enrichment of the *EVD/ATCOPIA93* retrotransposon, as originally reported. The full ECCsplorer pipeline was started with default settings and the trimming option enabled using the Nextera^TM^ adapter. A total of 501,558 circSeq PE reads and 143,372 control PE reads were mapped against the TAIR10 (The Arabidopsis Information Resource, http://www.arabidopsis.org) reference genome.

The ECCsplorer mapping module retrieved the well-known LTR retrotransposon *EVD/ATCOPIA93* as eccDNA candidate with the highest probability (Fig. 2a, orange) as well as other potential eccDNA candidates, e.g. originating from the ribosomal genes (Fig. 2a, blue, Supp. Tab. 4). In detail, the mapping module reported 13 highly confident candidate regions and 673 candidates with low confidence. The candidate region with the highest enrichment score of 56 was 5,332 bp long and annotated as *EVD* by the pipeline’s BLASTn analysis – in line with our expectation. We conclude that the ECCsplorer’s mapping module produces results that compare well with the original findings and that it is able to detect active retrotransposons from current datasets.

For the comparative read clustering, equal read counts of circSeq and control data were used (i.e. 96,016 PE reads per dataset). The clustering module identified 41.04 % of the clustered circSeq and 1.41 % of the control data reads as potential eccDNA candidates, respectively (39,402/1,371 reads from circSeq/control), pointing to a 28.7× overall increase (Fig. 2b). Four of the largest clusters containing about 10.3 % of all reads in candidate clusters (4,058/2 reads from circSeq/control), were reported as Ty1_copia/Ale, and correspond to the *EVD/ATCOPIA93* reference (Fig. 2b, orange circles, Supp. Tab. 5). We conclude that the eccDNA candidates can be readily detected with the clustering module as well as with the mapping module.

To be able to report highly confident eccDNA candidates, the pipeline compares the outputs of the mapping and clustering strategies. Here, the comparative module links eight candidate regions from the mapping module (Fig. 2a) to eleven clusters from the clustering module (Fig. 2b) with the best results for the eccDNA candidate *EVD/ATCOPIA93* (Fig 2c, Supp. Tab. 6). After removing duplicates, this leads to three target retrotransposon regions for potential experimental verification. These results are in line with the original study (Lanciano *et al*., 2017), and lead us to highlight the ECCsplorer’s potential for the fast and reliable identification of retrotransposon mobilization.

#### 3.1.3 Case study III: detection of (genic) eccDNAs using circSeq data from healthy humans (*H. sapiens*)

Typically, eccDNAs in *H. sapiens* are associated with cancer and other diseases. However, also in healthy tissues eccDNAs may arise (Møller *et al*., 2018): The study by Møller *et al.* generated multiple circSeq datasets from healthy blood and muscle tissues. They detected about 100,000 unique eccDNAs including genic eccDNAs. As circSeq enrichment data and the reference genome sequence were available, we re-analyzed these data using the ECCsplorer’s mapping module (`running mode: map`). As the original publication reported high eccDNA candidate density on chromosome 16 of the hg38 assembly (The 1000 Genomes Project Consortium *et al*., 2015), we used this chromosome as reference, along with the corresponding mRNA database as annotation database (UCSC Genome Browser, https://genome.ucsc.edu/).

The mapping module detected eccDNA candidates with high (n = 1) and low confidence (n = 840) on chr16 (Fig. 2d). The highly confident candidate region was 22,772 bp long and was located on the distal end of chromosome 16 (Fig. 2d, light blue, arrowed). It contains several gene annotations with the highest BLAST score observed for the gene mitofusin (MFN1, Supp. Tab. 8). Due to the arch-like distribution of the read coverage over the whole chromosome the candidate region with high confidence only showed an enrichment score of 0.16 as it is calculated globally. Although the ECCsplorer detected only a single eccDNA candidate region with high confidence, our manual analysis of regions with lower confidence showed also promising results. In total, 29 of those low-confidence candidate regions were supported by Møller et al.’s approach (Møller *et al*., 2018; Supp. Info. 3).

A main difficulty for this specific read dataset was the high background noise, presumably from linear DNAs that remained after incomplete exonuclease treatments. Nevertheless, the ECCsplorer pipeline was able to detect eccDNA candidates from genic regions, confirming the presence of eccDNAs in healthy *H. sapiens.*

#### 3.1.4 Case study IV: detection of eccDNAs absent from the reference genome using circSeq data from *B. vulgaris*

To test whether our ECCsplorer pipeline is able to detect eccDNAs absent from reference genome assemblies, we queried sugar beet (*B. vulgaris*), a non-model organism. The *B. vulgaris* genome harbors small extrachromosomal circular stretches of mitochondrial DNAs (Munk Hansen and Marcker, 1984) that are absent from the published reference assembly (Dohm *et al*., 2014; Funk *et al*., 2018). We generated low-coverage circSeq data (0.1×) from inflorescences of *B. vulgaris* after experimental enrichment of the eccDNA fraction according to Lanciano *et al.* (2017) and Diaz-Lara *et al.* (2016). As control, we used publicly available data from the same genotype (KWS2320, see data availability).

To test the ECCsplorer’s usability for the detection of extrachromosomal mitochondrial DNAs absent from the reference genome, we used the comparative read clustering as embedded in the clustering module (`running mode: clu`). For the clustering, 322,580 PE reads (161,290 PE reads per dataset) were used, respectively. The clustering module revealed twelve candidate clusters (combined in one supercluster) containing 30,792 reads (30,636/156 in circSeq/control). All were clearly enriched (Fig. 2e, orange circles, Supp. Tab. 9) and of mitochondrial origin. A manual BLAST assigned all twelve clusters to the *B. vulgaris*-typical mitochondrial minicircles, termed a, d and pO (Munk Hansen and Marcker, 1984).

These results clearly show that the ECCsplorer is capable of reference-free eccDNA detection at low sequencing coverages. We further want to highlight that already existing sequencing runs can be used for the comparative clustering analysis. Ideally, however, we recommend preparing enriched and non-enriched DNA from the same samples.

In contrast to other eccDNA detection methods, the ECCsplorer can be used without a high-quality reference genome sequence and is therefore much less vulnerable to assembly errors. This makes the ECCsplorer the current method of choice when working with non-model organisms.

### 3.2 Comparison with existing tools

Although the amount of available circSeq data and the interest in eccDNAs has been growing lately there is yet no standardized way of analyzing such data. To date, only very few attempts for software solutions are available and currently no solution is able to address different approaches in a single tool. Most of the available tools are aimed at a very specific use-case and are not applicable for non-model organisms at all.

To our knowledge there are currently three software solutions to detect eccDNAs. Circle-Map (Prada-Luengo *et al*., 2019) is a realigning-based pipeline to detect eccDNA from circSeq datasets already mapped to a reference genome sequence. Two further tools are available, though their very different application ranges prevent a direct comparison. Whereas the AmpliconArchitect (Deshpande *et al*., 2019) aims very specifically at detecting eccDNAs from *H. sapiens* cancer tissue, the CIDER-Seq (Mehta *et al*., 2020) approach relies on long-read (Pac-Bio/SMRT) sequences, only.

To evaluate the performance of our ECCsplorer pipeline, we compared its output with published results and those that we retrieved after re-running Circle-Map – to our knowledge the only tool directly comparable to the ECCsplorer. For the comparison, the datasets analyzed with the ECCsplorer (Fig. 2a/d) were reanalyzed with Circle-Map and bwa-mem (Li and Durbin, 2009) as mapping tool. In addition, we compare the results obtained from both tools with the originally published results. As Circle-Map relies on a gold-standard reference genome, only *A. thaliana* and *H. sapiens* data are suitable as input. To our knowledge, non-model organisms that lack high-quality draft genomes are only analyzable with our ECCsplorer pipeline when using short-read sequencing data at low coverage (< 0.2×).

First, we compared results of both tools and the originally published data from the enrichment of eccDNA in *A. thaliana* (Lanciano *et al*., 2017). In summary, the ECCsplorer detected eight eccDNA candidate regions and 673 regions with low confidence (Fig. 3a). Three of the eight candidates were annotated as *EVD/ATCOPIA93* (Fig 3a, first skyline plot, orange peaks), an LTR retrotransposon previously reported as transpositionally active (Mirouze *et al*., 2009). The five remaining candidates appeared to be organellar DNA or tandem repeats. The original publication (Lanciano *et al*., 2017) also reported eight candidate regions with three of them being *EVD/COPIA93* (Fig 4a, third skyline plot, orange peaks). None of the other candidates were originally annotated as organellar DNA as those reads had been filtered out before analysis. Re-analysis with the Circle-Map tool reported 1292 candidate regions, whereas regions greater than 50 kb have been manually filtered out as they were very likely false-positives or overlapped other candidates (Fig. 3a, second skyline plot, blue peaks).

**Figure 3:**
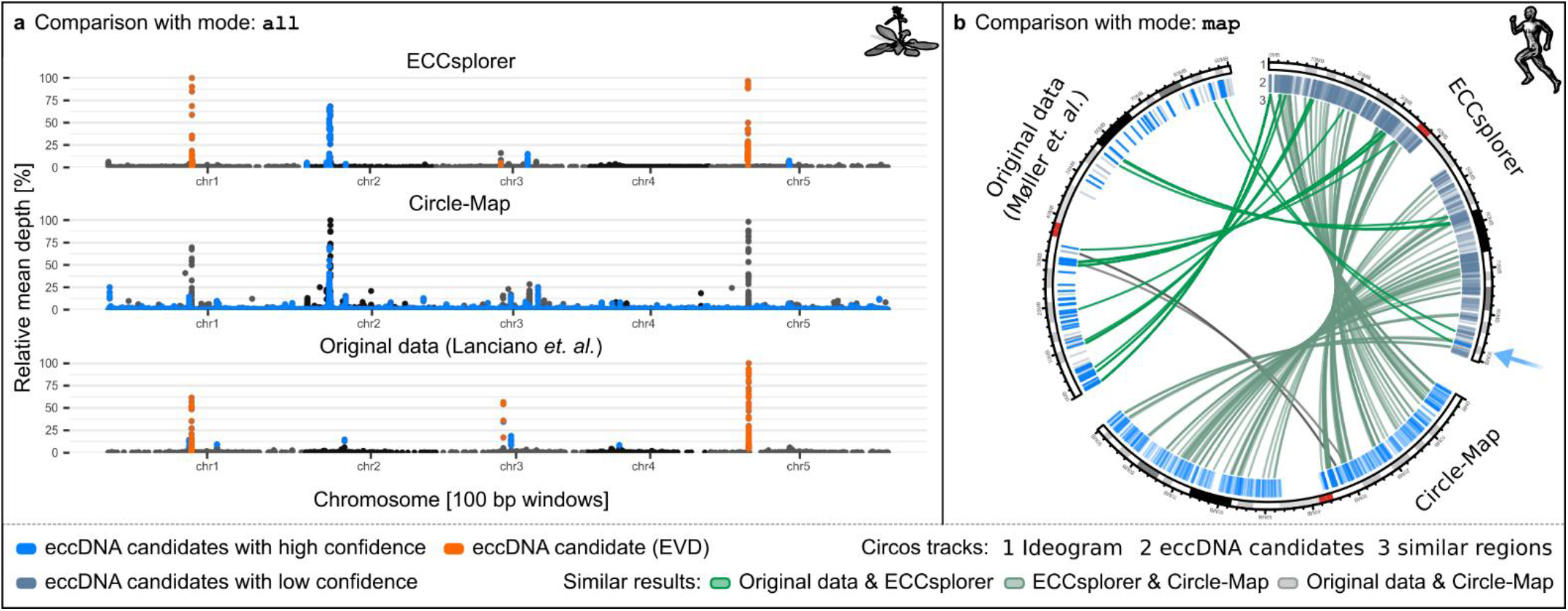
Comparison of the results from ECCsplorer pipeline (described here) with the Circle-Map (Prada-Luengo *et al*., 2019) and the originally published eccDNA candidates from *A. thaliana* (Lanciano *et al*., 2017) and *H. sapiens* (Møller *et al*., 2018). We reanalyzed data provided in Lanciano *et al.* (2017) and Møller *et al.* (2016) using the two available eccDNA tools, our ECCsplorer and Circle-Map. (**a**) Outputs using *A. thaliana* data from the different mapping analysis with the key eccDNA candidate *EVD/ATCOPIA93* (orange) and further candidate regions (blue) (maximal coverage depth of each dataset corresponds to 100 % relative depth). (**b**) Comparison of the different mapping analysis using *H. sapiens* data. The blue arrow marks one candidate with genic content from the ECCsplorer output.

The ECCsplorer output and the original data shared three high-confidence candidate regions with all of them being *EVD/ATCOPIA93* (Supp. Tab. 7). Four additional originally reported regions occurred in the low confidence output of the ECCsplorer pipeline. The ECCsplorer output and the Circle-Map output share three candidates as well, but none of them was *EVD/ATCOPIA93*. The Circle-Map output and the original data share five candidate regions, but again none of them being *EVD/ATCOPIA93*. There was no candidate region found by all three approaches (Supp. Tab. 7). This comparison demonstrates that the ECCsplorer pipeline provides more accurate results than the Circle-Map tool using default settings and considering the originally published results (Lanciano *et al*., 2017) as ground truth with experimental validation. Quite surprising was the absence of an *EVD/ATCOPIA93* eccDNA candidate in the Circle-Map results despite the large number of output candidates. This makes the ECCsplorer pipeline the currently best available option to detect active LTR retrotransposons from circSeq data in a unified available manner.

Second, we compared the results of ECCsplorer and Circle-Map with the originally published results of the *H. sapiens* circSeq study focusing on candidates on chromosome 16 from the hg38 assembly (Fig. 3b). Chromosome 16 was reported by the original study (Møller *et al*., 2018) to have a high per Mb eccDNA count and is also one of the shorter *H. sapiens* chromosomes allowing all tools to run on a desktop-grade computer. The ECCsplorer pipeline found one candidate region with high and 840 with low confidence using default setting (Fig. 3b), respectively. The original study (Møller *et al*., 2018) reported 70 highly and 75 lowly confident candidate regions. Re-analysis with the Circle-Map tool found 493 candidates using recommended settings, and manual filtering of candidates greater than 50 kb. The outputs of all approaches were compared with BEDtools intersect (for details see Supp. methods). The ECCsplorer pipeline had 29 results in common with the originally published data and 82 overlaps to candidates reported by the Circle-Map tool (Fig 3b, Supp. Info. 3). Curiously, only two similar candidates were shared between the original data and the Circle-Map results. No candidates were shared across all approaches. This demonstrates the complexities of the detection of eccDNA candidates from circSeq data and the need of a unified, and reproducible software solution. The ECCsplorer pipeline shared candidates with both other approaches, while these shared almost none among themselves. This finding makes the ECCsplorer currently the best starting point for the analysis of eccDNAs. This is especially true in the light that the published candidates from *H. sapiens* (Møller *et al*., 2018) have not entirely been experimentally validated. Additionally, our findings demonstrate that an approach based on mapping only may result in many false-positive eccDNA candidates. Therefore, we recommend running the ECCsplorer pipeline with all implemented approaches, including the mapping, clustering, and comparative modules.

## 4 Conclusion

Due to the wide relevance in medicine and breeding, eccDNA analyses are gaining popularity. Nevertheless, the detection of eccDNAs from next-generation enrichment sequencing is far from being a solved problem, and there is a high demand for new tools. In particular, organisms lacking gold-standard genome assemblies are virtually impossible to analyze to date. To meet this need, we introduce our ECCsplorer pipeline that modularly combines reference-free and reference-guided procedures to identify eccDNA candidates from rolling-circle amplification protocols such as circSeq and mobilome-seq. Using simulated data (case study I) as well as real world data (case studies II-IV) from various organisms, we were able to demonstrate the ECCsplorer pipeline’s functionality, showcase its wide applicability, and highlight its advantages over existing tools. Finally, the usage of a bioinformatics pipeline for the analysis of data examining a fairly new research field allows advanced comparability between different studies and ensures their reproducibility.

## Acknowledgments

The authors are grateful for support and mentorship from the late Thomas Schmidt. Furthermore, we thank Stefan Wanke for supporting our research initiative. We thank the laboratory of Marie Mirouze (IRD, Marseille) for kindly providing the mobilome-Seq protocol and the publicly available circSeq (mobilome-Seq) data shown in Fig. 2 and 3. Similarly, we acknowledge the laboratory of Brigitte Regenberg (University of Copenhagen) for the publicly available circSeq data shown in Fig. 2, 3. The KWS Saat SE & Co. KGaA are acknowledged for providing plant material. We thank N. Schmidt for beta testing of the software, and G. Menzel for his support.

## Author contributions

LM developed ECCsplorer, and wrote the manuscript. KMS provided input in the development of ECCsplorer. BW and TH conceived the study and designed the experiments. All authors were involved in discussion of the manuscript, in editing, and approved the final manuscript.

## Supplementary information

Supplementary Information 1: Supplementary Methods. Detailed methods for circSeq data generation and running the ECCsplorer pipeline, including all parameters and commands.

Supplementary Information 2: Supplementary Tables 1-9.

Supplementary Information 3: Supplementary Data. All data used for plots in Figures 1–3.

