## Supplementary Information 1: Supplementary Methods for "ECCsplorer: a pipeline to detect extrachromosomal circular DNA (eccDNA) from next-generation sequencing data"

### 1 Methods

#### 1.1 Plant material and DNA extraction

*Beta vulgaris ssp. vulgaris* genotype KWS2320 was grown under greenhouse conditions by KWS Saat SE & Co. KGaA. Freshly harvested inflorescences were shipped on ice, stored at 4 °C and whole genomic DNA was extracted within a week. For one pooled plant sample, whole genomic DNA was extracted using the plant DNeasy mini kit (Qiagen) according to the manufacturer's instructions.

#### 1.2 Extrachromosomal circular DNA enrichment

For the removal of large genomic linear fragments 3 µg of genomic DNA was purified using a QIAquick PCR purification kit (Qiagen) according to the manufacturer's instructions. Remaining linear DNA from 25 µl of the purification product was further removed using the Plasmid-Safe™ ATP-dependent DNase (Epicentre) exonuclease according to the manufacturer's instructions, except that the incubation at 37 °C was elongated to 17 h. The DNA sample was cleaned up using a standard ethanol/glycogen precipitation protocol by adding 0.1 volume of sodium acetate (3M), 2.5 volume ethanol and 1 µl glycogen (Beckman Coulter). The precipitated circular DNA was enriched by random rolling circle amplification (rRCA) using the Illustra™ TempliPhi™ amplification kit (GE Healthcare). Therefore, the DNA pellet from the precipitation was directly resuspended in 5 µl of the TempliPhi™ sample buffer. The reaction was performed according to the manufacturer's instructions, except that the incubation at 28 °C was performed for 65 h. For the determination of the DNA concentration, 1 µl of the enriched DNA sample was purified using the GeneJet DNA clean up kit (Thermo Fisher Scientific) according to the manufacturer's instructions. Then, the DNA concentration was measured optically using a NanoDrop™

photometer (Thermo Fisher Scientific). The remaining sample was diluted to 1 ng/µl for library preparation and sequencing.

#### 1.3 Library preparation and sequencing

Library preparation using the Nextera™ library kit and sequencing was performed through the commercial provider Macrogen (Macrogen Inc., Seoul, Korea, www.macrogen.com). Sequencing was performed using the HiSeq X Ten platform (Illumina) targeting 20 Gb of 2 x 250 nucleotide paired-end reads. A mapping to the reference genome sequence EL10 (Funk *et al.*, 2018) using bowtie2 (Langmead and Salzberg, 2012) with default settings revealed an average insert size of around 650 bp.

#### 1.4 Data analysis and running of the ECCsplorer pipeline

In order to illustrate and test the ECCsplorer's operating principle, we created semi-artificial test datasets, available on https://github.com/crimBubble/ECCsplorer/tree/master/testdata. These contain a region (0.6 Mb) from the *B. vulgaris* reference genome sequence as the control dataset. To simulate retrotransposon enrichment typical for some eccDNAs, we added multiple concatenated *Beetle7* copies with LTR/LTR and solo-LTR junctions to obtain circSeq data. For the creation of the artificial circSeq and the control datasets (25,000 spots each with 2 x 200 bp) we ran dwgsim (https://github.com/nh13/DWGSIM) with default settings. The ECCsplorer pipeline was run with the trimming option enabled and the read count option set to auto.

```
$ python3 ECCsplorer.py aDNA_R1.fastq aDNA_R2.fastq gDNA_R1.fastq
gDNA_R2.fastq -trm tru2 -d RefSeq_DB.fasta -ref RefGenSeq.fasta -
cnt 3000 –dsa circsim –dsb control
```

For the validation of the ECCsplorer's functionality, we profited from some of the first eccDNA enrichment studies available (Lanciano *et al.*, 2017; Møller *et al.*, 2018). CircSeq data from *A.*

*thaliana* (ERR1830501 and corresponding control datasets ERR1830499), and similar data from *H. sapiens* muscle tissue (SRR6315430) were used, respectively. As references for the mapping module, the current reference genome assemblies were used, namely the *A. thaliana* TAIR10 (The Arabidopsis Information Resource, http://www.arabidopsis.org) and the *H. sapiens* hg38 (UCSC Genome Browser, https://genome.ucsc.edu/). Additionally, the corresponding gene and mRNA databases were retrieved as annotation databases. For both datasets, the ECCsplorer pipeline was run with the trimming option enabled and for the *H. sapiens* dataset the window size parameter has been increased to 250 bp.

```
$ python3 ECCsplorer.py ERR1830501_1.fastq.gz ERR1830501_2.fastq.gz ERR1830499_1.fastq.gz ERR1830499_2.fastq.gz -trm nex -d Araport11_genes.201606.cds.fasta ATCOPIA93.fasta -ref TAIR10.fa -dsa epi12 -dsb cntrl
```

```
$ python3 ECCsplorer.py SRR6315430_1.fastq.gz SRR6315430_2.fastq.gz -trm tru2 -d GRCh38_rna.fna -ref hg38_chr16.fa –win 250
```

For the reference-free detection eccDNA-enriched *B. vulgaris* reads were used (ERR6004146), respectively. As control, WGA reads from the same genotype, used for generation of the published genome assembly (Dohm *et al.*, 2014) (SRR869631), were retrieved. The ECCsplorer was run with the trimming option enabled, read count set to auto (using a genome coverage of 0.1×) with an estimated genome size of 750 MB and with otherwise standard parameters.

```
$ python3 ECCsplorer.py ERR6004146_1.fastq.gz ERR6004146_2.fastq.gz SRR6315430_1.fastq.gz SRR869631_2.fastq.gz -trm nex –cnt auto –rgs 750000000
```

#### 1.5 Comparison between the CircleMap and the ECCsplorer softwares

To enable a software comparison between the ECCsplorer and Circle-Map (Prada-Luengo *et al.*, 2019), circSeq data from *A. thaliana* and *H. sapiens* were mapped against the corresponding reference genomes. To achieve a typical Circle-Map output, the by the authors provided tutorial instructions for mapping and the identification of circular DNAs were followed (https://github.com/iprada/Circle-Map/wiki/Tutorial:-Identification-of-circular-DNA-using-Circle-Map-Realign).

BED files were retrieved from the outputs of the ECCsplorer and the Circle-Map tools, and from the provided material of the original studies (Lanciano *et al.*, 2017; Møller *et al.*, 2018). These were compared using BEDtools intersect (-u option) (Quinlan, 2014). The results have been visualized using the R packages ggplot2 (Wickham, 2016) (https://ggplot2.tidyverse.org/) and circlize (Gu *et al.*, 2014) (https://jokergoo.github.io/circlize/).

#### 1.6 Hardware and required software

All ECCsplorer pipeline runs were performed on a Unix machine (Ubuntu 16.04 LTS, i7 6-gen with 64 GB). The ECCsplorer pipeline is implemented in Python 3 (3.5 or higher) with Biopython (Cock *et al.*, 2009), Scipy (Virtanen *et al.*, 2020) and pyRserve (Heinkel, 2017) and R (R Core Team, 2013) with ggplot2 (Wickham, 2016), ggrepel (Slowikowski *et al.*, 2019), gridExtra (Auguie and Antonov, 2017) and dplyr (Wickham *et al.*, 2019) and 3$^{rd}$ party tools Blast+ (Camacho *et al.*, 2009), Trimmomatic (Bolger *et al.*, 2014) (optional but recommended), seqtk (Li *et al.*, 2013) (optional but recommended for better performance), segemehl (Hoffmann *et al.*, 2014), SAMtools (Li *et al.*, 2009), BEDtools (Quinlan, 2014), RepeatExplorer2 (Novák *et al.*, 2010, 2020).

### 2 Additional references (methods)


Auguie, B. and Antonov, A. (2017) gridExtra: miscellaneous functions for “grid” graphics.

Bolger, A.M. *et al.* (2014) Trimmomatic: a flexible trimmer for Illumina sequence data. *Bioinformatics*, **30**, 2114–2120.

Camacho, C. *et al.* (2009) BLAST+: architecture and applications. *BMC Bioinformatics*, **10**, 421.

Cock, P.J.A. *et al.* (2009) Biopython: freely available Python tools for computational molecular biology and bioinformatics. *Bioinformatics*, **25**, 1422–1423.

Dohm, J.C. *et al.* (2014) The genome of the recently domesticated crop plant sugar beet (*Beta vulgaris*). *Nature*, **505**, 546–549.

Funk, A. *et al.* (2018) Nucleotide-binding resistance gene signatures in sugar beet, insights from a new reference genome. *The Plant Journal*.

Gu, Z. *et al.* (2014) circlize implements and enhances circular visualization in R. *Bioinformatics*, **30**, 2811–2812.

Heinkel, R. (2017) pyRserve: A Python client to remotely access the R statistic package via network.

Hoffmann, S. *et al.* (2014) A multi-split mapping algorithm for circular RNA, splicing, trans-splicing and fusion detection. *Genome Biology*, **15**, R34–R50.

Lanciano, S. *et al.* (2017) Sequencing the extrachromosomal circular mobilome reveals retrotransposon activity in plants. *PLoS Genetics*, **13**, e1006630–e1006650.

Langmead, B. and Salzberg,S.L. (2012) Fast gapped-read alignment with Bowtie 2. *Nature Methods*, **9**, 357–359.

Li, H. *et al.* (2013) Seqtk: a fast and lightweight tool for processing FASTA or FASTQ sequences.

Li, H. *et al.* (2009) The sequence alignment/map format and SAMtools. *Bioinformatics*, **25**, 2078–2079.

Møller, H.D. *et al.* (2018) Circular DNA elements of chromosomal origin are common in healthy human somatic tissue. *Nat Commun*, **9**, 1069–1081.

Novák, P. *et al.* (2020) Global analysis of repetitive DNA from unassembled sequence reads using RepeatExplorer2. *Nature Protocols*, **15**, 3745–3776.

Novák, P. *et al.* (2010) Graph-based clustering and characterization of repetitive sequences in next-generation sequencing data. *BMC Bioinformatics*, **11**, 378–390.

Prada-Luengo, I. *et al.* (2019) Sensitive detection of circular DNAs at single-nucleotide resolution using guided realignment of partially aligned reads. *BMC Bioinformatics*, **20**, 663.

Quinlan, A.R. (2014) BEDTools: The Swiss-Army Tool for Genome Feature Analysis. *Curr Protoc Bioinformatics*, **47**, 11.12.1-34.

R Core Team (2013) R: A language and environment for statistical computing R Foundation for Statistical Computing, Vienna, Austria.

Slowikowski, K. *et al.* (2019) ggrepel: automatically position non-overlapping text labels with ‘ggplot2’.

Virtanen, P. *et al.* (2020) SciPy 1.0: fundamental algorithms for scientific computing in Python. *Nature Methods*, **17**, 261–272.

Wickham, H. *et al.* (2019) dplyr: a grammar of data manipulaton. R package version 0.8. 0.1.

Wickham, H. (2016) ggplot2: Elegant Graphics for Data Analysis Springer.
