## Supplementary Information 2: Supplementary Tables 1-9 for "ECCsplorer: a pipeline to detect extrachromosomal circular DNA (eccDNA) from next-generation sequencing data"

#### Supplementary Table 1 (semi-artificial data, `running mode: all > mapping module`)

Supplementary Table 1: Top results from mapping module using test data (circsim and control). Using this dataset, the one artificially created eccDNA candidate was detected.

| Candidate | Enrichment score | Length [bp] | Position | Best hit annotation |
| --- | --- | --- | --- | --- |
| eccCand_001 | 46.5 | 6695 | chr3:86641-93336 | Beetle-7#LTR/Gypsy/chromovirus/CRM@Beta_vulgaris |

#### Supplementary Table 2 (semi-artificial data, `running mode: all > clustering module`)

Supplementary Table 2: Top results from clustering module using test data (circsim and control). Using this dataset, the one artificially created eccDNA candidate was detected as one supercluster.

| Cluster | Supercluster | Size [reads] | epiRIL proportion | Best hit annotation |
|---|---|---|---|---|
| CL1 | 1 | 401 | 0.97 | All/repeat/mobile_element/Class_I/LTR/Ty3_gypsy/chromovirus/CRM |
| CL2 | 1 | 370 | 0.97 | All/repeat/mobile_element/Class_I/LTR/Ty3_gypsy/chromovirus/CRM |
| CL3 | 1 | 330 | 0.979 | All/repeat/mobile_element/Class_I/LTR/Ty3_gypsy/chromovirus/CRM |
| CL4 | 1 | 321 | 0.984 | All/repeat/mobile_element/Class_I/LTR/Ty3_gypsy/chromovirus/CRM |
| CL5 | 1 | 291 | 0.973 | All/repeat/mobile_element/Class_I/LTR/Ty3_gypsy/chromovirus/CRM |
| CL6 | 1 | 280 | 0.979 | All/repeat/mobile_element/Class_I/LTR/Ty3_gypsy/chromovirus/CRM |
| CL7 | 1 | 260 | 0.992 | All/repeat/mobile_element/Class_I/LTR/Ty3_gypsy/chromovirus/CRM |
| CL8 | 1 | 254 | 0.992 | All/repeat/mobile_element/Class_I/LTR/Ty3_gypsy/chromovirus/CRM |
| CL9 | 1 | 215 | 0.967 | All/repeat/mobile_element/Class_I/LTR/Ty3_gypsy/chromovirus/CRM |
| CL10 | 1 | 209 | 0.981 | All/repeat/mobile_element/Class_I/LTR/Ty3_gypsy/chromovirus/CRM |
| CL11 | 1 | 206 | 0.981 | All/repeat/mobile_element/Class_I/LTR/Ty3_gypsy/chromovirus/CRM |
| CL12 | 1 | 172 | 0.977 | All/repeat/mobile_element/Class_I/LTR/Ty3_gypsy/chromovirus/CRM |
| CL13 | 1 | 166 | 0.976 | All/repeat/mobile_element/Class_I/LTR/Ty3_gypsy/chromovirus/CRM |

#### Supplementary Table 3 (semi-artificial data, `running mode: all > comparative module`)

Supplementary Table 3: Top results from comparative module using test data (circsim and control). Using this dataset, the one artificially created eccDNA candidate was detected.

| Candidate | Enrichment score | Best hit annotation (mapping module) | Associated clusters | epiRIL proportion (per cluster) | Best hit annotation (clustering module) |
|---|---|---|---|---|---|
| eccCand_001 | 46.5 | Beetle-7#LTR/Gypsy/chromovirus/CRM@Beta_vulgaris | CL1 | 0.97 | All/repeat/mobile_element/Class_I/LTR/Ty3_gypsy/chromovirus/CRM |
| | | | CL2 | 0.97 | |
| | | | CL3 | 0.979 | |
| | | | CL4 | 0.984 | |
| | | | CL5 | 0.973 | |
| | | | CL6 | 0.979 | |
| | | | CL7 | 0.992 | |
| | | | CL8 | 0.992 | |
| | | | CL9 | 0.967 | |
| | | | CL10 | 0.981 | |
| | | | CL11 | 0.981 | |
| | | | CL12 | 0.977 | |
| | | | CL13 | 0.976 | |

#### Supplementary Table 4 (*A. thaliana* data, `running mode: all > mapping module`)

Supplementary Table 4: Top results from mapping module using *A. thaliana* data (epi12 and WT). Using this data set, 13 eccDNA candidate regions were detected.

<table>
<tr><th>Candidate</th><th>Enrichment score</th><th>Length [bp]</th><th>Position</th><th>Best hit annotation</th></tr>
<tr><td>eccCand_001</td><td>56</td><td>5332</td><td>chr5:5629975-5635307</td><td>EVD AT5TE20395|+|5629978|5635310|ATCOPIA93|LTR/Copia|5333</td></tr>
<tr><td>eccCand_002</td><td>28</td><td>6649</td><td>chr1:12753744-12760393</td><td>EVD AT5TE20395|+|5629978|5635310|ATCOPIA93|LTR/Copia|5333</td></tr>
<tr><td>eccCand_003</td><td>12</td><td>424</td><td>chr5:11813591-11814015</td><td></td></tr>
<tr><td>eccCand_004</td><td>10</td><td>127</td><td>chr3:10182070-10182197</td><td>EVD AT5TE20395|+|5629978|5635310|ATCOPIA93|LTR/Copia|5333</td></tr>
<tr><td>eccCand_005</td><td>7</td><td>744</td><td>chr2:5829018-5829762</td><td>AT2G13895.1</td></tr>
<tr><td>eccCand_006</td><td>4.5</td><td>8863</td><td>chr2:3500250-3509113</td><td>150_gi|26556996|ref|NC_001284.2|#organelle/mitochondria</td></tr>
<tr><td>eccCand_007</td><td>2</td><td>34116</td><td>chr2:3346113-3380229</td><td>150_gi|26556996|ref|NC_001284.2|#organelle/mitochondria</td></tr>
<tr><td>eccCand_008</td><td>2</td><td>12029</td><td>chr5:11838404-11850433</td><td></td></tr>
<tr><td>eccCand_009</td><td>1.33</td><td>2971</td><td>chr2:1622-4593</td><td>23_gi|604854740|gb|KJ507198.1|#45S_rDNA/18S_rDNA</td></tr>
<tr><td>eccCand_010</td><td>1.33</td><td>943</td><td>chr2:2937853-2938796</td><td>443_gi|7525012|ref|NC_000932.1|#organelle/plastid</td></tr>
<tr><td>eccCand_011</td><td>1</td><td>3446</td><td>chr2:6785-10231</td><td>24_25S_PS_CL2Contig6_rc/3985-7377#45S_rDNA/25S_rDNA</td></tr>
<tr><td>eccCand_012</td><td>1</td><td>4015</td><td>chr2:3268139-3272154</td><td>150_gi|26556996|ref|NC_001284.2|#organelle/mitochondria</td></tr>
<tr><td>eccCand_013</td><td>1</td><td>25814</td><td>chr3:14191918-14217732</td><td>24_25S_PS_CL2Contig6_rc/3985-7377#45S_rDNA/25S_rDNA</td></tr>
</table>

#### Supplementary Table 5 (semi-artificial data, `running mode: all > clustering module`)

Supplementary Table 5: Top 20 results from clustering module using *A. thaliana* data (epi12 and WT). Using this data set, multiple eccDNA candidate regions were detected by the clustering approach.

| Cluster | Super cluster | Size [reads] | epiRIL proportion | Best hit annotation |
|---|---|---|---|---|
| CL1 | 2 | 4060 | 1.0 | All/repeat/mobile_element/Class_I/LTR/Ty1_copia/Ale |
| CL2 | 3 | 3373 | 0.857 | All/organelle/mitochondria |
| CL6 | 3 | 2530 | 0.804 | All/organelle/mitochondria |
| CL10 | 10 | 2362 | 1.0 | All |
| CL14 | 2 | 2212 | 0.999 | All/repeat/mobile_element/Class_I/LTR/Ty1_copia/Ale |
| CL16 | 2 | 1923 | 1.0 | All/repeat/mobile_element/Class_I/LTR/Ty1_copia/Ale |
| CL17 | 8 | 1796 | 1.0 | All |
| CL18 | 3 | 1629 | 0.878 | All/organelle/mitochondria |
| CL19 | 9 | 1480 | 1.0 | All |
| CL23 | 14 | 1345 | 1.0 | All |
| CL26 | 8 | 1303 | 1.0 | All |
| CL32 | 18 | 1138 | 0.998 | All |
| CL33 | 9 | 1129 | 1.0 | All |
| CL37 | 20 | 943 | 1.0 | All |
| CL42 | 23 | 903 | 0.998 | All |
| CL49 | 28 | 750 | 1.0 | All |
| CL56 | 35 | 628 | 0.994 | All/repeat |
| CL61 | 39 | 572 | 1.0 | All |
| CL63 | 2 | 544 | 0.996 | All/repeat/mobile_element/Class_I/LTR/Ty1_copia/Ale |
| CL78 | 53 | 401 | 1.0 | All |

### Supplementary Table 6 (semi-artificial data, `running mode: all > comparative module`)

Supplementary Table 6: Top results from comparative module using *A. thaliana* data (epi12 and WT). Using this data set, eight eccDNA candidate regions were detected across both approaches (mapping and clustering) and are considered eccDNA candidates with high confidence.

| Candidate | Enrichment score | Best hit annotation (mapping module) | Associated clusters | epiRIL proportion (per cluster) | Best hit annotation (clustering module) |
|---|---|---|---|---|---|
| eccCand_001 | 56 | EVD AT5TE20395\|+\|5629978\|5635310\|<br>ATCOPIA93\|LTR/Copia\|5333 | CL1<br>CL14<br>CL16<br>CL63 | 1.0<br>0.999<br>1.0<br>0.996 | All/repeat/mobile_element/<br>Class_I/LTR/Ty1_copia/Ale |
| eccCand_002 | 28 | EVD AT5TE20395\|+\|5629978\|5635310\|<br>ATCOPIA93\|LTR/Copia\|5333 | CL1<br>CL14<br>CL16<br>CL63 | 1.0<br>0.999<br>1.0<br>0.996 | All/repeat/mobile_element/<br>Class_I/LTR/Ty1_copia/Ale |
| eccCand_004 | 10 | EVD AT5TE20395\|+\|5629978\|5635310\|<br>ATCOPIA93\|LTR/Copia\|5333 | CL1 | 1.0 | All/repeat/mobile_element/<br>Class_I/LTR/Ty1_copia/Ale |
| eccCand_006 | 4.5 | 150_gi\|26556996\|ref\|NC_001284.2\|<br>#organelle/mitochondria | CL2<br>CL6<br>CL18<br>CL415 | 0.857<br>0.804<br>0.878<br>0.926 | All/organelle/mitochondria |
| eccCand_007 | 2 | 150_gi\|26556996\|ref\|NC_001284.2\|<br>#organelle/mitochondria | CL2<br>CL6<br>CL18<br>CL415 | 0.857<br>0.804<br>0.878<br>0.926 | All/organelle/mitochondria |
| eccCand_011 | 1 | 24_25S_PS_CL2Contig6_rc/3985-7377<br>#45S_rDNA/25S_rDNA | CL288<br>CL405 | 0.956<br>0.862 | All/repeat/rDNA/45S_rDNA/<br>25S_rDNA |
| eccCand_012 | 1 | 150_gi\|26556996\|ref\|NC_001284.2\|<br>#organelle/mitochondria | CL89 | 0.825 | All |
| eccCand_013 | 1 | 24_25S_PS_CL2Contig6_rc/3985-7377<br>#45S_rDNA/25S_rDNA | CL288<br>CL405 | 0.956<br>0.862 | All/repeat/rDNA/45S_rDNA/<br>25S_rDNA |

### Supplementary Table 7 (Comparison of ECCsplorer, Circle-Map, and originally published candidates)

Supplementary Table 7: Overlapping candidate regions using *A. thaliana* data (epi12 and WT) from each mapping analysis (**ECC**splorer pipeline, **C**ircle-**M**ap, and originally published data by **Lan**ciano et al. (2017)) examined using BEDtools.

| Chromosome | Start | End | Quality | Method | Chromosome | Start | End | Quality | Method |
|---|---|---|---|---|---|---|---|---|---|
| chr1 | 12753744 | 12760393 | hconf | ECC | chr1 | 12754300 | 12760400 | hconf | Lan |
| chr2 | 3268139 | 3272154 | hconf | ECC | chr2 | 3268140 | 3270259 | --- | CM |
| chr2 | 3268139 | 3272154 | hconf | ECC | chr2 | 3270663 | 3273277 | --- | CM |
| chr2 | 3346113 | 3380229 | hconf | ECC | chr2 | 3344596 | 3346513 | --- | CM |
| chr2 | 3346113 | 3380229 | hconf | ECC | chr2 | 3349795 | 3350330 | --- | CM |
| chr2 | 3346113 | 3380229 | hconf | ECC | chr2 | 3350818 | 3353174 | --- | CM |
| chr2 | 3346113 | 3380229 | hconf | ECC | chr2 | 3355398 | 3356724 | --- | CM |
| chr2 | 3346113 | 3380229 | hconf | ECC | chr2 | 3358494 | 3359166 | --- | CM |
| chr2 | 3346113 | 3380229 | hconf | ECC | chr2 | 3359287 | 3378942 | --- | CM |
| chr2 | 3500250 | 3509113 | hconf | ECC | chr2 | 3499864 | 3501375 | --- | CM |
| chr3 | 10182070 | 10182197 | hconf | ECC | chr3 | 10181900 | 10182400 | hconf | Lan |
| chr5 | 5629975 | 5635307 | hconf | ECC | chr5 | 5630000 | 5635300 | hconf | Lan |
| chr1 | 12362558 | 12363626 | --- | CM | chr1 | 12362700 | 12363200 | hconf | Lan |
| chr1 | 16582926 | 16583270 | --- | CM | chr1 | 16583000 | 16583200 | hconf | Lan |
| chr2 | 5829019 | 5829758 | --- | CM | chr2 | 5829200 | 5829400 | hconf | Lan |
| chr3 | 11357284 | 11358090 | --- | CM | chr3 | 11357300 | 11357900 | hconf | Lan |
| chr4 | 4594263 | 4594725 | --- | CM | chr4 | 4594400 | 4594600 | hconf | Lan |

#### Supplementary Table 8 (semi-artificial data, `running mode: map > mapping module`)

Supplementary Table 8: Top results from mapping module using *H. sapiens* data (circSeq). Only one eccDNA candidate region was detected with high confidence.

Additionally, 841 eccDNA candidate region were detected with low confidence.

| Candidate | Enrichment score | Length [bp] | Position | Best hit annotation |
|---|---|---|---|---|
| eccCand_001 | 0.16 | 22722 | chr16:87568894-87591666 | MFN1 mitofusin 1 [NM_033540.3] |

#### Supplementary Table 9 (semi-artificial data, `running mode: clu > clustering module`)

Supplementary Table 9: Top results from clustering module using *B. vulgaris* data (circSeq and WGA). Cluster contigs were annotated by a nucleotide similarity search (online BLAST) against the NCBI database with default settings (organism ~ viridiplantae).

| Cluster | Super cluster | Size [reads] | epiRIL proportion | Best hit annotation | NCBI Blast best hit annotation |
|---|---|---|---|---|---|
| *CL3* | 1 | 5593 | 0.996 | All/organelle/mitochondria | mitochondrial minicircle a/d [X04983.1, X04984.1] |
| *CL5* | 1 | 3510 | 0.998 | All/organelle/mitochondria | mitochondrial minicircle d [X04984.1] |
| *CL6* | 1 | 3292 | 0.994 | All/organelle/mitochondria | mitochondrial minicircle a [X04983.1] |
| *CL10* | 1 | 2691 | 0.989 | All/organelle/mitochondria | mitochondrial minicircle pO [X00641.1] |
| *CL11* | 1 | 2633 | 0.994 | All/organelle/mitochondria | mitochondrial minicircle a [X04983.1] |
| *CL13* | 1 | 2343 | 0.997 | All/organelle/mitochondria | mitochondrial minicircle d [X04984.1] |
| *CL15* | 1 | 2140 | 0.999 | All/organelle/mitochondria | mitochondrial minicircle d [X04984.1] |
| *CL16* | 1 | 2099 | 0.994 | All/organelle/mitochondria | mitochondrial minicircle a [X04983.1] |
| *CL17* | 1 | 1958 | 0.996 | All/organelle/mitochondria | mitochondrial minicircle d [X04984.1] |
| *CL18* | 1 | 1954 | 0.994 | All/organelle/mitochondria | mitochondrial minicircle a [X04983.1] |
| *CL21* | 1 | 1763 | 0.997 | All/organelle/mitochondria | mitochondrial minicircle d [X04984.1] |
| *CL38* | 1 | 816 | 0.987 | All/organelle/mitochondria | mitochondrial minicircle d/pO [X04984.1, X00641.1] |
